# IMMUNOGENICITY AND SAFETY OF CHICKEN EMBRYO FIBROBLAST ADAPTED FOWLPOX VACCINE

**DOI:** 10.1101/2025.02.12.637982

**Authors:** Uchenna Somtochukwu Okafor, Stephen Israel, Clement Adebajo Meseko, Oyeduntan Adejoju Adediran, Oluyemi Rotimi Ogunmolawa, Anthony B. Darang, Dauda Garba Bwala

## Abstract

**Background:** The use of chorioallantoic membranes (CAM) route for the propagation of fowlpox virus (FPV) is the method applied in the production of potent and effective fowlpox vaccine at the National Veterinary Research Institute (NVRI), Vom. However, production insufficiency has led to inadequate supply of vaccine to meet farmer’s demands. This has left large number of susceptible and unvaccinated flocks with outbreaks of the disease being reported in Nigerian.

**Aim:** This study adapted FPV to Chicken embryo fibroblast (CEF) cell culture and assessed its immunogenicity and safety of the resulting vaccine in comparison to a locally produced CAM-based FPV vaccine.

**Methods:** The FPV were propagated on CAM and CEF of 9 to 12 day old developing chicken embryos. The vaccine harvests were subjected to quality checks, titrated, and used in vaccinating experimental birds via the wing web to monitor for takes, seroconversion, and safety.

**Results:** The FPV was successful adapted to CEF as demonstrated by the attainment of 80 to 90% cytopathic effect within 71 to 115 hours in the different passages. Both the CEF and CAM-adapted vaccine harvests were highly replicative producing titres of above 10^6.0^/ml TCID_50,_ but higher titres of 10^8.25^/ml TCID_50_ was recorded in the CEF-adapted vaccines. Experimentally vaccinated birds showed 100% takes within 3 to 4 days post-vaccination, with no adverse effects recorded.

**Conclusion:** This research reported successful development and adaptation of FPV to CEF that was safe and immunogenic, with the potential for its use in the production of self-sufficiency vaccine for fowlpox prevention and control in poultry.

## INTRODUCTION

Fowl pox is known to be a global poultry disease caused by a DNA virus belonging to the genus *Avipoxvirus*, under the family *Poxviridae*. The growth of proliferative lesions and scabs on the skin, as well as diphtheritic lesions in the upper portions of the respiratory and digestive tracts, are the hallmarks of this gradual spreading illness (Tripathy & Reed, 2020; WOAH, 2023). Although fowlpox affects both domestic and wild birds, it primarily affects chickens and turkeys, but cases of infection have been reported in other avian species such as ducks, geese, pheasants, quail, canaries, pigeons, and hawks (Bwala *et al*., 2015; Giotis and Skinner, 2018; 2021) with variable incidences depending on the climatic conditions, management and hygiene or the practice of regular vaccination (WOAH, 2023).

Although fowlpox is listed as a disease of economic importance by the World Animal Health Organization (WOAH), it only temporarily lowers egg production and slows the growth of young birds. Mortality rate and economic losses in younger birds suffering from the diphtheritic form could be higher than in the cutaneous form, sometimes reaching up to 50%. Mortality from the cutaneous form can also be high especially when lesions develop around the eyes which affects the bird’s vision and ability to feed (Tripathy & Reed, 2020). Fowlpox has been controlled in most developed countries but remained a problem in many developing countries including Nigeria largely due to poor hygiene, contaminated instruments/surfaces, and vector transmission via biting insects such as mosquitoes and other flies, and poultry red mites (*Dermanyssus gallinae*) (Tripathy & Reed, 2020). Nigeria has a large prevalence of fowlpox, which primarily affects backyard and free-range flocks but can also infect flocks of chickens and turkeys that are intensively raised (Adene and Fatumbi, 2004; Meseko *et al*., 2012; Meseko *et al.,* 2017).

There are no definitive treatments for fowl pox, thus control and prevention mainly depend on maintaining good hygiene and timely vaccination. Vaccination remains a potent and the most effective way of preventing and controlling fowlpox in domestic birds. Thus, a wide variety of commercially available vaccines have been tested in birds of different age ranges with varying degrees of success and different levels of protection (Woodward and Tudor, 1973; Baxi and Oberoi, 1999). These fowlpox vaccines which are mostly live vaccines are produced either by inoculation onto chorioallantoic membranes (CAM) of 9- to 12-day-old developing chicken embryos or cell cultures of avian origin. A number of cell cultures have previously been employed in the propagation of fowlpox virus, including chicken embryo kidney cells, primary chicken embryo fibroblasts (CEF), chicken embryo dermis cells, or the permanent quail cell line QT-35 (Schnitzlein *et al*., 1988). The use of the CAM route for the propagation of the fowlpox virus (WOAH, 2023) has been the method applied in the production of the fowlpox vaccine at the National Veterinary Research Institute (NVRI), Vom, and the vaccine has been effective in the prevention and control of fowlpox in poultry flocks. However, the quantity produced is grossly inadequate to meet the demand of farmers and are thus supplemented by import. The imported vaccines are still inadequate and are highly exorbitant and out of the reach of poor small scale poultry farmers. This leads to poor vaccine coverage with large number of susceptible unvaccinated flocks; thus outbreaks and infections of flocks are regularly being reported in Nigeria (Odoya *et al*., 2006).

Fowlpox vaccine in most countries is propagated on CAM, but in modern veterinary vaccinology, vaccines derived from cell culture are now widely used (Yusifova, 2021; WOAH, 2023). The immunogenicity, efficacy, and safety of cell culture-adapted fowlpox in comparison to fowlpox vaccines of other origins have been investigated, yielding diverse results (Baxi and Oberoi, 1999; El-Mahdy and Mikheal, 2014; Wambura and Mzula, 2017; Radwan and Mikhael, 2020). However, the overall reports have consistently indicated positive outcomes for the cell culture-adapted fowlpox vaccines. These vaccines, whether administered alone or in combination with other vaccines, have demonstrated effectiveness and safety with several biological advantages of the cell-culture-adapted vaccines, including product uniformity, homogeneity, and stringent bacterial and fungal sterility (El-Zein, 1974).

Barhouna and Hanson (1986) have reported the embryo’s fibroblast as one of the cells that can be used for the multiplication of fowlpox viruses, but Vero cells have also been used to propagate this virus. Several studies have evaluated chicken embryo fibroblast cell culture adapted fowl pox vaccines and were reported to be safe with a high ability to induce protective immune response in vaccinated birds (Baxi and Oberoi, 1999; Yusifova, 2018; Sarma *et al*. 2019; Khalili *et al*., 2020). In addition, the use of cell culture-adapted vaccines for fowlpox has been reported to be more economical and productive compared to CAM-adapted vaccines whose production is more time-consuming, thus numerous producers began manufacturing vaccines using cell systems (Yusifova, 2021). However, there are generally no information on either the production or study on cell culture-adapted fowlpox vaccines in Nigeria. Furthermore, in view of cases of reported outbreaks of fowlpox in vaccinated flocks, it has become imperative to develop vaccines against fowlpox that have strong immunogenic qualities (Yusifova, 2017; 2018; 2021). Thus, with a view to developing a more potent vaccines in enough quantity to meet the demand of the Nigerian poultry industry this study adapted fowlpox vaccine virus to CEF and assessed its safety and immunogenicity.

## MATERIALS AND METHODS

### Ethical Approval for the Experiment

Ethical approval for the use of sentinels in this experiment was procured from the NVRI Animal Use and Care Committee (AEC/02/141/23) before the experiment commenced.

### Propagation of vaccine virus

The vaccine virus was propagated both in chicken embryo fibroblast (CEF) and chorioallantoic membrane (CAM) of Specific Antibody Negative (SAN) eggs obtained from the National Veterinary Research Institute Vom poultry farm. The Primary fibroblast cells were prepared from 9-11-day-old chicken embryos. The chicken embryo fibroblast cell culture for the propagation of the vaccine virus was produced in-house and the working virus seed was inoculated in both suspension and on monolayer as described by Khalili and colleagues (2020). The CAM propagation of the vaccine virus was carried out by inoculating approximately 0.1 ml of the seed virus onto the CAMs of 9- to 12-day-old developing chicken embryos (WOAH, 2023).

### Master Seed Virus (MSV) Preparation of Virus

Two different commercial vaccine vials (BIOVAC and IZOVAC) were procured from a reputable vendor in Jos, both of which were live attenuated fowlpox vaccines propagated on CAM from SPF eggs. Each vial of 1000 doses of freeze-dried live vaccine were reconstituted in 2ml of sterile HMEM (Hanks Minimum Essential Medium), giving a titer of 10^3^ EID_50_/dose. The resulting constitute was used as the working seed for the vaccine production.

### Chicken embryo fibroblast (CEF) cell culture

Cell culture was done using primary fibroblast cells obtained from 9 to 12 days old chick embryo of SAN flock as described (Cotter *et al*., 2001; Cunningham, 1966; Hernandez *et al*., 2010). Briefly described, the limbs and heads of the harvested embryos were cut off and embryos macerated and washed several times with PBS (phosphate buffered saline) solution of pH 7.4 and two changes of trypsin solution. The macerated tissue pieces were further digested with THBSS (trypsin Hanks balanced salt solution) using magnetic stirrer and magnetic apparatus. The digestion process was repeated several times until the embryo was fully digested. The digestion process was halted by adding fetal bovine serum (FBS). The suspension containing the digested cells was then filtered through a coarse muslin gauze. The resulting filtrate was centrifuged at 2500 rpm for 20 minutes to precipitate cells. The resulting supernatant was decanted leaving the cells at the bottom. The sedimented cells were resuspended in HMEM in a 1: 350 dilutions. A liter of the medium was supplemented with 10% Newborn calf serum and 10mls of 1% penicillin-streptomycin to prevent infection. 5 ml of fibroblast cell suspensions were cultured in sterile 25ml-cell culture flasks for the production of monolayer fibroblast cells and incubated at 37^0^c. Both suspension and monolayer culture flasks were prepared for virus inoculation.

### Adaptation of FPV in CEF cell culture

The suspension CEF primary cells in HMEM media were inoculated with 0.2ml virus suspension and incubated at 37^0^C. For the production of Monolayer CEF, the primary cells in HMEM media were cultured and incubated for 12-24 hours at 37^0^C. After 70-80% confluency of the cells had been obtained, the media was poured out and 0.2ml of virus suspension was inoculated into mono-layered CEF cells and incubated at 37°C for 45mins - 1 hour for the adsorption of the virus to take place after which non-adsorbed virus was decanted and a maintenance medium (HMEM containing 0.1% antibiotics and 2% Newborn calf serum) was added, incubated at 37^0^C and observed daily for 3-5days until the cytopathic effect (CPE) reaches between 80 – 90% when the vaccines are frozen and harvested. Harvested vaccines were then subjected to quality checks including sterility and titer determination (Olfat, 2005; WOAH, 2023). The harvested vaccine was thereafter passaged in CEF cells for up to five passages.

### Vaccine Titration

The titers of the MSV and the Fowlpox virus vaccines were obtained by 2 methods: 50 per cent or median Embryo Infectious Dose (EID_50_) and TCID_50_ (Median Tissue Culture Infectious Dose) (Reed and Muench, 1938; Khalili *et al*., 2020; WOAH, 2023).

### Median Embryo Infectious Dose **-** EID_50_ of CAM-based vaccines

Specific antibody-negative (SAN) embryonated chicken eggs were inoculated via the CAM route with 0.2 ml of the ten-fold serial dilution of the virus suspension. Five (5) embryonated eggs (9-12 days old) were inoculated for each dilution (10^1^ – 10^10^). The embryonated eggs that were found dead within 24 hours post inoculations (PI) were not considered. The embryos that died after 24 hours and those that survived were examined on day 5 PI for evidence of viral activity/infection (presence of pock lesions or generalized thickening of CAM). The EID_50_ was calculated using the Reed and Muench method (1938).

### Median Tissue Culture Infectious Dose - TCID_50_ of CEF vaccines

Primary CEF cells were prepared in 24-well microplates, and virus dilution from 10^-1^ to 10^-10^ was prepared, 5 wells were allocated to each dilution. Equal volume (1ml) of media and virus suspension were added into each well, while one well remained as control. The plates were incubated (37°C; 5% CO_2_) and monitored daily for CPE and on the fifth day post inoculation. The TCID_50_/ml was calculated using the Spearman-Karber formula (M = xk + 1 / 2d-drl / n) (Spearman, 1908; K rber, 1931).

### Vaccine Compounding

Some portion of the harvests from the cell culture-adapted vaccine of the various passages were compounded with the requisite excipients which include peptone, gelatin and antibiotics. The resulting mixture was then dispensed into sterile vaccine vials, partially stoppered and freeze-dried. The freeze-dried vaccines were then subjected to quality assurance checks and titration

### Safety test

For the safety test, three groups of 5 birds each were vaccinated with CEF-adapted vaccine with half field dose (25μl), field dose (50μl), and 10 times (10x) field doses (500μl). CAM-based vaccine was administered to two groups of 5 birds each for field dose (50μl), and 10 times field dose (500μl). The CAM-based half-field dose group were not included due to shortage of space, coupled with the fact that data already exist where mostly field doses are administered to chickens in poultry farms. The CEF-adapted vaccine half-field dose was included in this study because it is a new experimental vaccine. A control group of 5 birds were maintained without vaccination. The groupings and vaccination schedule are outlined in Table 1.

**Table 1:** Experimental design showing the different groupings and their treatment/durations.

| Groups | Week |  |  |  |  |  |  |
| --- | --- | --- | --- | --- | --- | --- | --- |
|  | 1 | 2 | 3 | 4 | 5 | 6 | 7 |
|  |  |  | (7 DPV) | (14 DPV) | (21 DPV) | (28 DPV) | (35 DPV) |
| **A1**<br>(5 Birds) | Stabilize<br>& bleed | Vaccinate (CEF<br>vac) half dose | Bleed all<br>birds | Bleed all<br>birds | Bleed all<br>birds | Bleed all<br>birds | Bleed all<br>birds |
| **A2**<br>(5 Birds) | Stabilize<br>& bleed | Vaccinate (CEF<br>vac) std dose | Bleed all<br>birds | Bleed all<br>birds | Bleed all<br>birds | Bleed all<br>birds | Bleed all<br>birds |
| **A3**<br>(5 Birds) | Stabilize<br>& bleed | Vaccinate (CEF<br>vac) 10x std dose | Bleed all<br>birds | Bleed all<br>birds | Bleed all<br>birds | Bleed all<br>birds | Bleed all<br>birds |
| **B2**<br>(5 Birds) | Stabilize<br>& bleed | Vaccinate (CAM<br>vac) std dose | Bleed all<br>birds | Bleed all<br>birds | Bleed all<br>birds | Bleed all<br>birds | Bleed all<br>birds |
| **B3**<br>(5 Birds) | Stabilize<br>& bleed | Vaccinate (CAM<br>vac) 10x std dose | Bleed all<br>birds | Bleed all<br>birds | Bleed all<br>birds | Bleed all<br>birds | Bleed all<br>birds |
| **Control**<br>(5 Birds) | Stabilize<br>& bleed | No vaccination | Bleed all<br>birds | Bleed all<br>birds | Bleed all<br>birds | Bleed all<br>birds | Bleed all<br>birds |

**Key:** DPV = Days Post-vaccination, A1 (group that received CEF-adapted vaccine - half field dose); A2 (group that received CEF-adapted vaccine - field dose); A3 (group that received CEF- adapted vaccine - 10 times field dose); B2 (group that received CAM vaccine - field dose); B3 (group that received CAM vaccine - 10 times field dose); and Control (unvaccinated group).

The experimental birds were vaccinated by piercing the wing web with a two-pronged needle dipped into each vaccine type. The vaccinated chickens were then observed daily for 7–10 days for evidence of ‘takes’ (A ‘take’ consists of swelling of the skin or the formation of a scab at the site of vaccination which is considered evidence of successful vaccination), and adverse effects/clinical disease attributable to the vaccines. The birds were further monitored for 5 weeks and were bled weekly to check for seroconversion.

### Antibody Decay Monitoring

For this purpose, the 30 birds above were bled at regular intervals as outlined in Table 1. To monitor for the fowlpox-specific antibody seroconversion and the antibody profile, the weekly post-vaccination sera were stored for analysis at -20^O^C until analyzed by Enzyme-Linked Immunosorbent Assay (ELISA) (Chicken Fowlpox Virus Antibody (FPV-Ab) ELISA Kit) (Khalili *et al*., 2020). Due to the shortage of the ELISA plate, only a representative sample from each group was analyzed, while the remaining sera were also subjected to analysis by Agar gel immunodiffusion (AGID) test.

### Enzyme Linked Immunosorbent Assay (ELISA) Test

An FPV antibody ELISA kit (Shanghai Melson Medical) was used to analyze sera samples randomly selected from samples of each group of different doses and different weeks from the vaccinated and control groups according to the manufacturer’s instruction and the World Organization for Animal Health (WOAH, 2023). Results were read at 450nm wavelength and recorded. The cutoff OD (optical density) was calculated according to manufacturer’s instructions and recorded. All sera samples could not be assayed because the ELISA kit only came with one 96-well plate, and thus could not accommodate all the samples.

### Agar Gel Immunodiffusion (AGID) Test

For this purpose, known FPV antigen-containing (Charles River laboratories) control samples were used for FPV antibody detection in selected sera samples. Agarose gel of 1.5% was prepared and poured into petri dishes. When the gel was set, a gel puncher was used to punch the plates, creating a central well and 6 peripheral wells. The known antigen samples were placed in the central well, while the serum samples to be analyzed were placed in the peripheral wells and labelled appropriately. The plates were then incubated at 37 ^0^c and the results of precipitation lines were read within 24-72 hrs. Post-vaccination serum samples that could not be included in the ELISA due to limited test wells were analyzed with AGID. A total of 64 serum samples were analyzed via this technique. The assay was carried out as described (WOAH, 2023).

## RESULTS

### Development of primary cell line and vaccine virus propagation

The development of primary CEF cell line that was used for the propagation of the vaccine virus and also for titration of the harvests and the freeze-dried vaccines obtained a cell count of 125 x 10^4^ CEF cells/ml; 75cm**^2^** Corning**^®^**cell culture flasks (plug seal cap type) were used for virus propagation. The virus propagation and passages using cell culture were performed in both suspension and monolayer, and freezing and harvest of products in all 5 passages were only done after 80 – 90% CPE were observed in each passage. The CPE recorded, the lack of CPE in the control and the different timelines for the termination of incubation of the different passages on the attainment of 80 – 90% CPE are indicated in Figure 1, Figure 2 and Table 2 respectively. The thickening of the CAM resulting from the propagation of the virus on CAM is presented in Figure 3.

**Figure 1:**
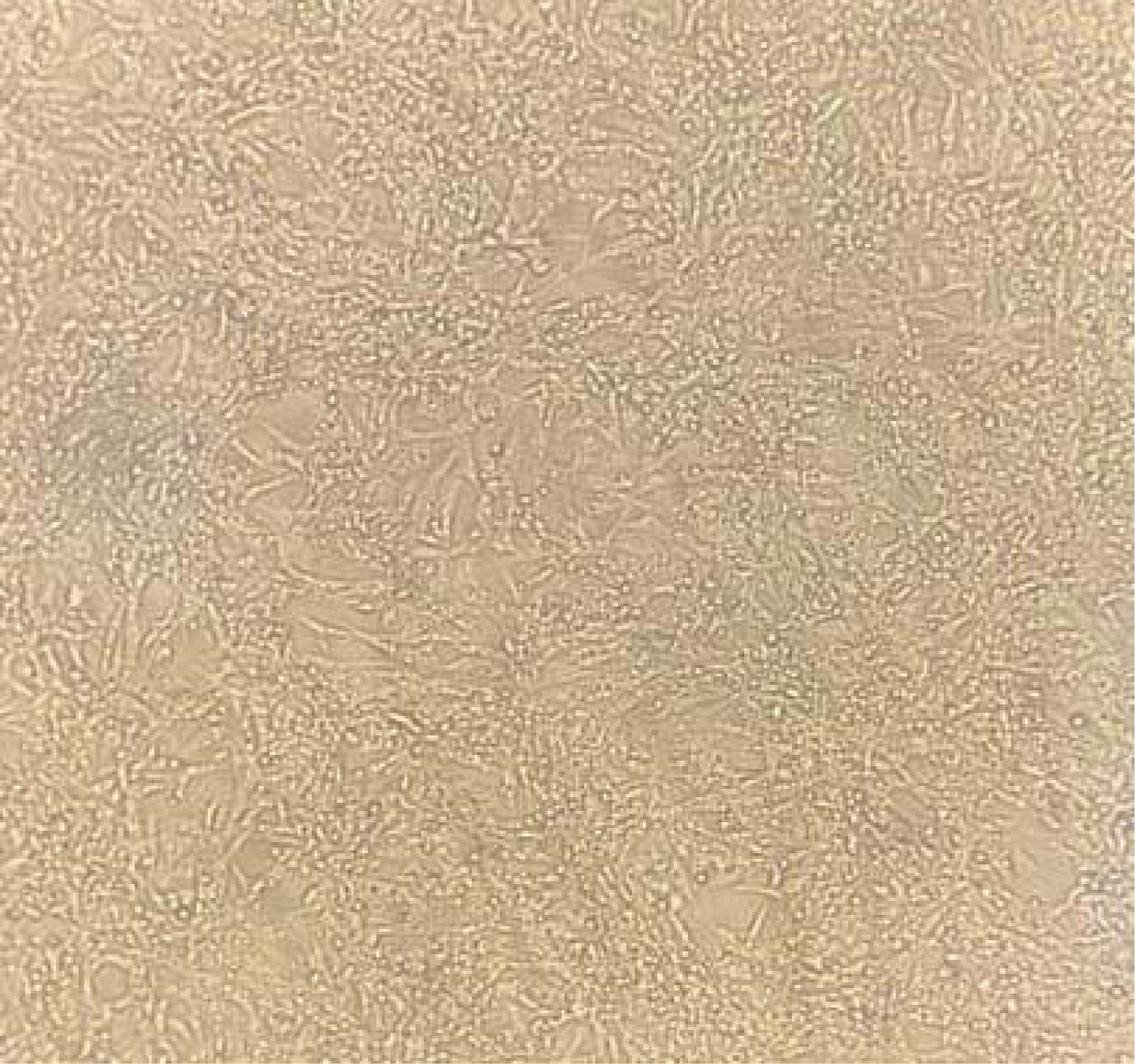
CEF culture inoculated with fowlpox virus showing CPE (rounding of cells and formation of syncytia) (passage 2 cells).

**Figure 2:**
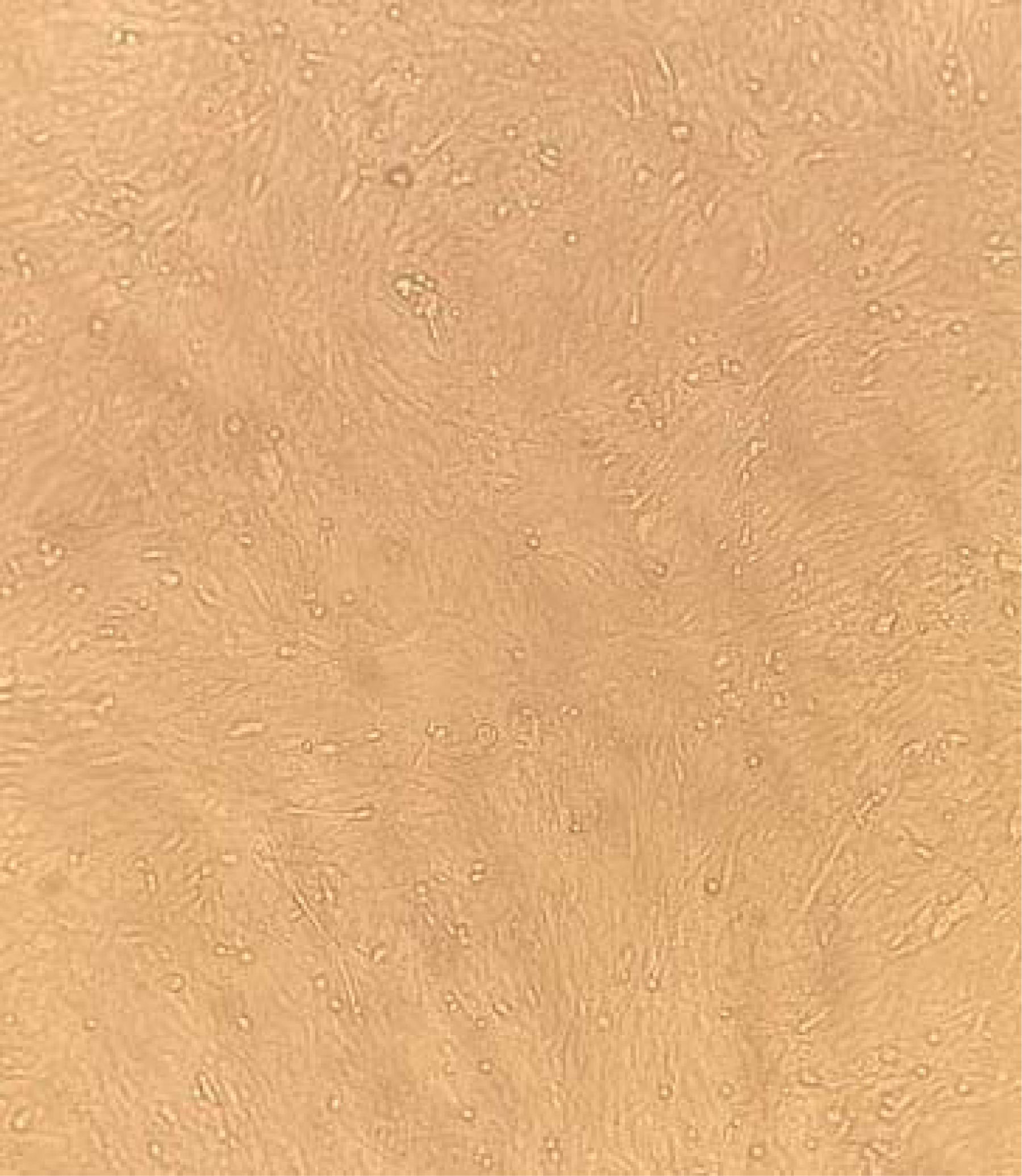
Uninoculated Control CEF culture without CPE at the termination of incubation.

**Figure 3:**
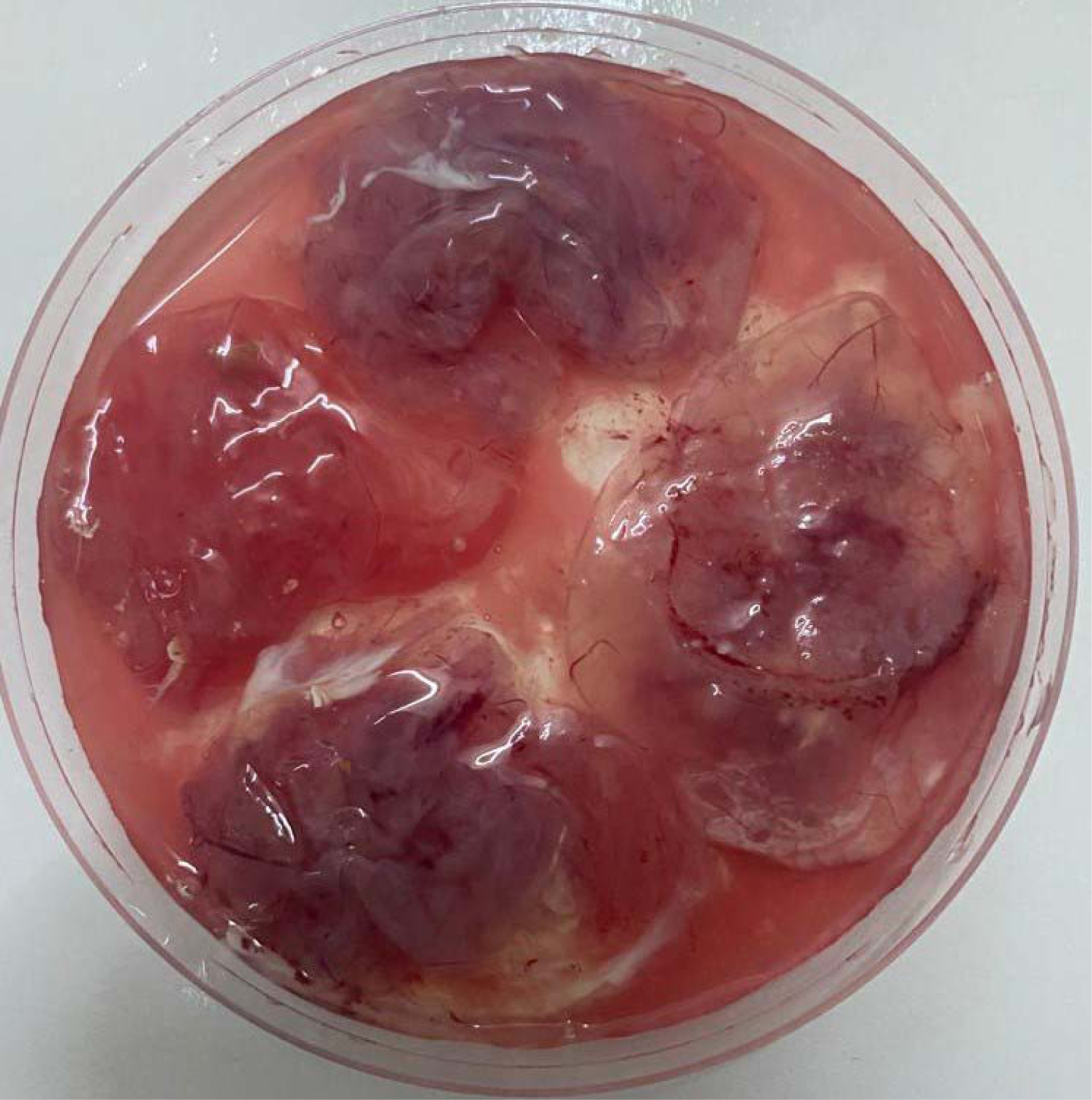
Thickened CAM due to pocks formation on FPV-infected CAM (5 days PI).

**Table 2:** Termination time of incubation of virus propagation in CEF cell cultures after attaining 80%-90% CPE.

| Cell Culture Type | Cell Culture Passages |  |  |  |  |
| --- | --- | --- | --- | --- | --- |
|  | 1 | 2 | 3 | 4 | 5 |
| Suspension | 115hrs | 85hrs 22mins | 71hrs 14mins | 93hrs | 90hrs |
| Monolayer | 85hrs 5mins | 45hrs | 48hrs | 69hrs | 60hrs |

Key: CPE = Cytopathic Effect, CEF = Chicken Embryo Fibroblast Cell Culture

### Immunogenicity of the CAM and CEF-adapted vaccines

The vaccine virus seeds: BIOVAC (Seed 1) and IZOVAC (Seed 2) inoculated and propagated on CAM and the various passaged in CEF cell culture as well as the freeze-dried harvests (seed 2) titers were determined and presented in Table 3.

**Table 3:** TCID_50_ titers of different passages of fowlpox vaccine harvest and freeze Dried vaccines grown in CEF cell culture.

| Vaccine Type | Culture type | Culture Passages |  |  |  |  |
| --- | --- | --- | --- | --- | --- | --- |
|  |  | 1 | 2 | 3 | 4 | 5 |
| Vaccine Seed 1 | Suspension | 10<sup>6.5</sup>/ml | 10<sup>6.5</sup>/ml | 10<sup>6.5</sup>/ml | DC | DC |
|  | Monolayer | 10<sup>6.5</sup>/ml | 10<sup>6.5</sup>/ml | 10<sup>6.7</sup>/ml | DC | DC |
| Vaccine Seed 2 | Suspension | 10<sup>6.7</sup>/ml | 10<sup>6.7</sup>/ml | 10<sup>7.2</sup>/ml | 10<sup>8.0</sup>/ml | 10<sup>8.2</sup>/ml |
|  | Monolayer | 10<sup>7.5</sup>/ml | 10<sup>7.0</sup>/ml | 10<sup>7.2</sup>/ml | 10<sup>8.2</sup>/ml | 10<sup>7.7</sup>/ml |
| Freeze Dried Vaccine<br>(Seed 2) | Suspension | 10<sup>6.5</sup>/ml | 10<sup>6.5</sup>/ml | 10<sup>6.5</sup>/ml | 10<sup>7.0</sup>/ml | 10<sup>7.2</sup>/ml |
|  | Monolayer | 10<sup>7.5</sup>/ml | 10<sup>6.5</sup>/ml | 10<sup>6.5</sup>/ml | 10<sup>7.2</sup>/ml | 10<sup>7.7</sup>/ml |

Keys: Vaccine Seed 1 – Biovac; Vaccine Seed 2 – Izovac; DC = Discontinued because of the low titers of harvests from passages 1 – 3. TCID<sub>50</sub> = Median Tissue Culture Infectious Dose; CEF = Chicken Embryo Fibroblast Cell culture.

The ELISA results of the 3-4 representative sera samples from each group from chickens vaccinated with the varying doses of CEF-adapted and CAM vaccines over the 5-week observation period are presented in Table 4. The 82 samples randomly selected and analyzed, were all positive with optical density (OD) readings above the cutoff value (0.216). The negative control gave optical density (OD) readings of 0.066 which is below the critical cutoff OD figures of 0.216 as provided by the manufacturer of the kit (values lower than 0.216 were regarded as negative results). On the other hand, the positive control gave OD readings of 3.310 and 3.251 for the ELISA test which confirmed their positivity (values higher than or equal to 0.216 were considered to be positive for FPV antibody). Chickens vaccinated with CEF-adapted vaccine generally have higher OD values signifying the presence of higher antibody titer as shown in Table 4.

**Table 4:**
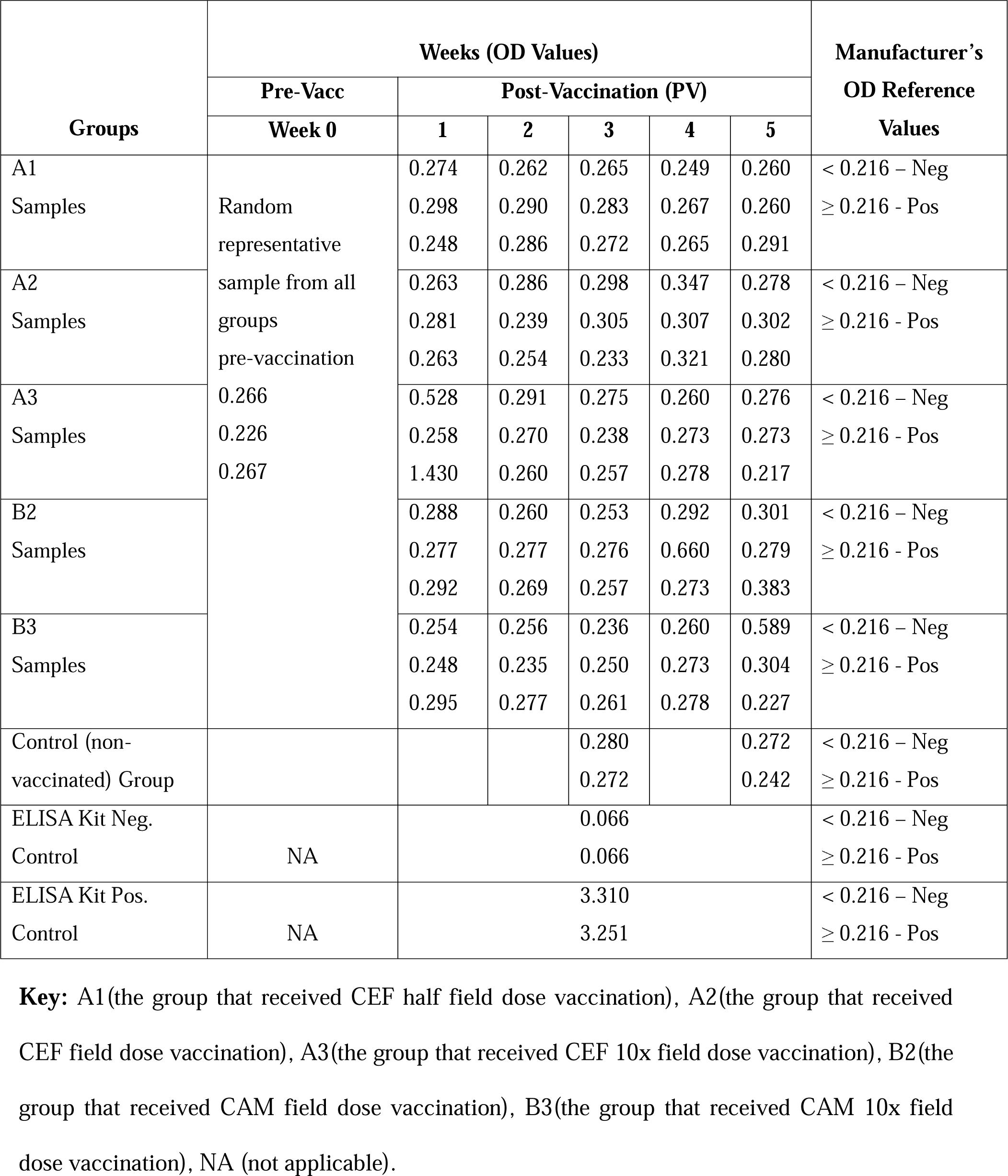
ELISA result showing the optical density (OD) readings of FPV antibody values post-vaccination for 5 weeks.

Furthermore, the sera samples that could not be analyzed using ELISA due to the shortage of space in the one ELISA Microtiter plate that accompanied the kit, were analyzed using AGID. Of the total 62 samples analyzed using AGID, 18 (29%) were positive for FPV antibody as presented in Table 5, The result showed all the groups having positive results except the unvaccinated control group and pre-vaccination baseline samples. The treatment group A2 which received a field dose of CEF-adapted vaccine had the highest percentage positivity while the treatment group A3 which received 10x field dose of CEF-adapted vaccine did not produce the highest percentage positivity.

**Table 5:**
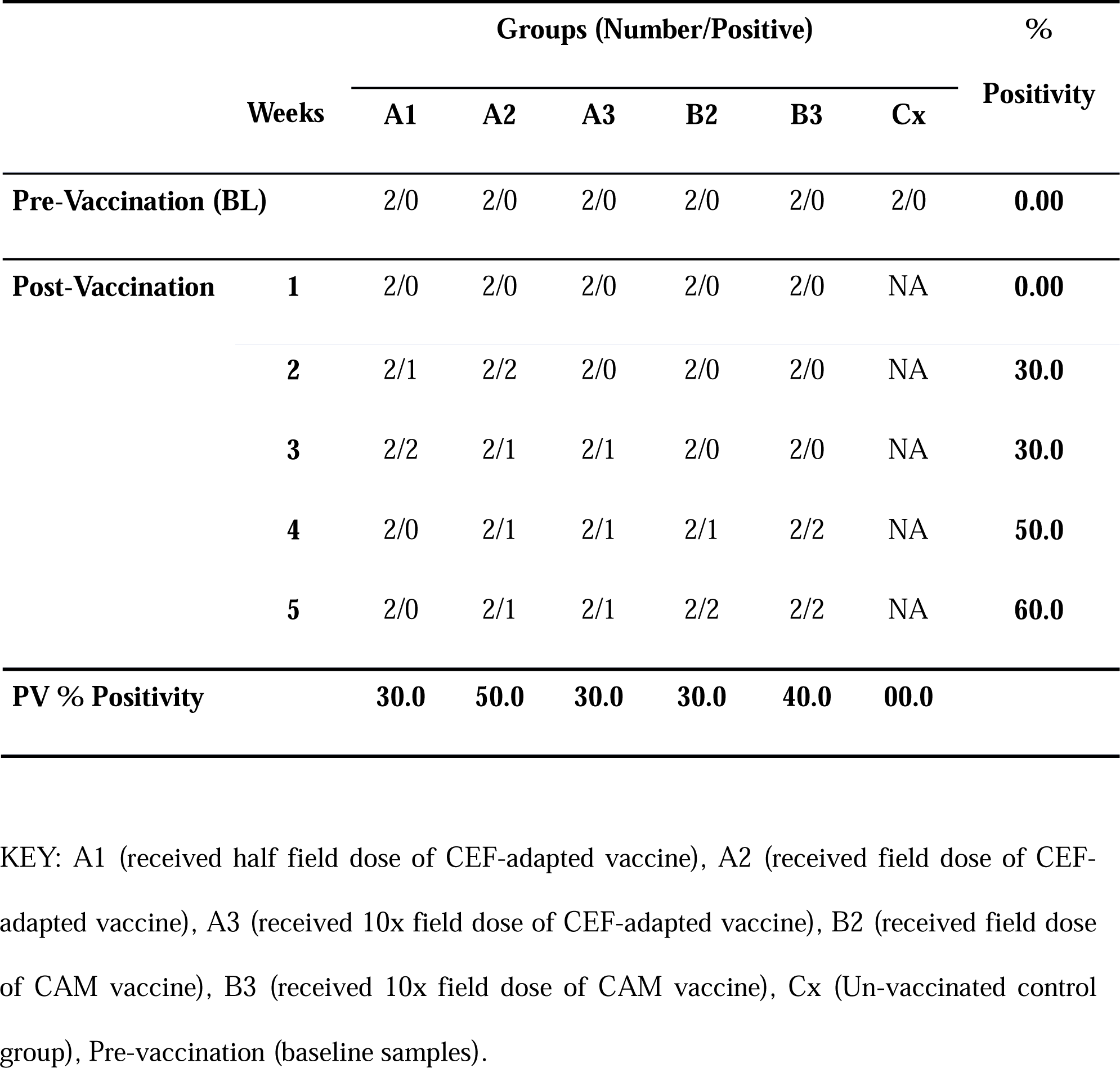
Pre- and post-vaccination sera samples analyzed by AGID.

### Safety of CAM and CEF-adapted vaccines

The broiler chicks vaccinated with the varying doses of CEF-adapted and CAM vaccines were monitored daily for “takes” (swelling and/or scab formation at site of injection) and other untoward or adverse signs attributable to the vaccine. “Takes” were observed in all the vaccinated birds within three to four days post vaccination as shown in Figure 4. The unvaccinated control groups did not show any evidence of “takes”. In addition, the safety of the vaccines was confirmed across the varying doses as there were no adverse effects or clinical signs observed during the 5weeks monitoring period post vaccination.

**Figure 4:**
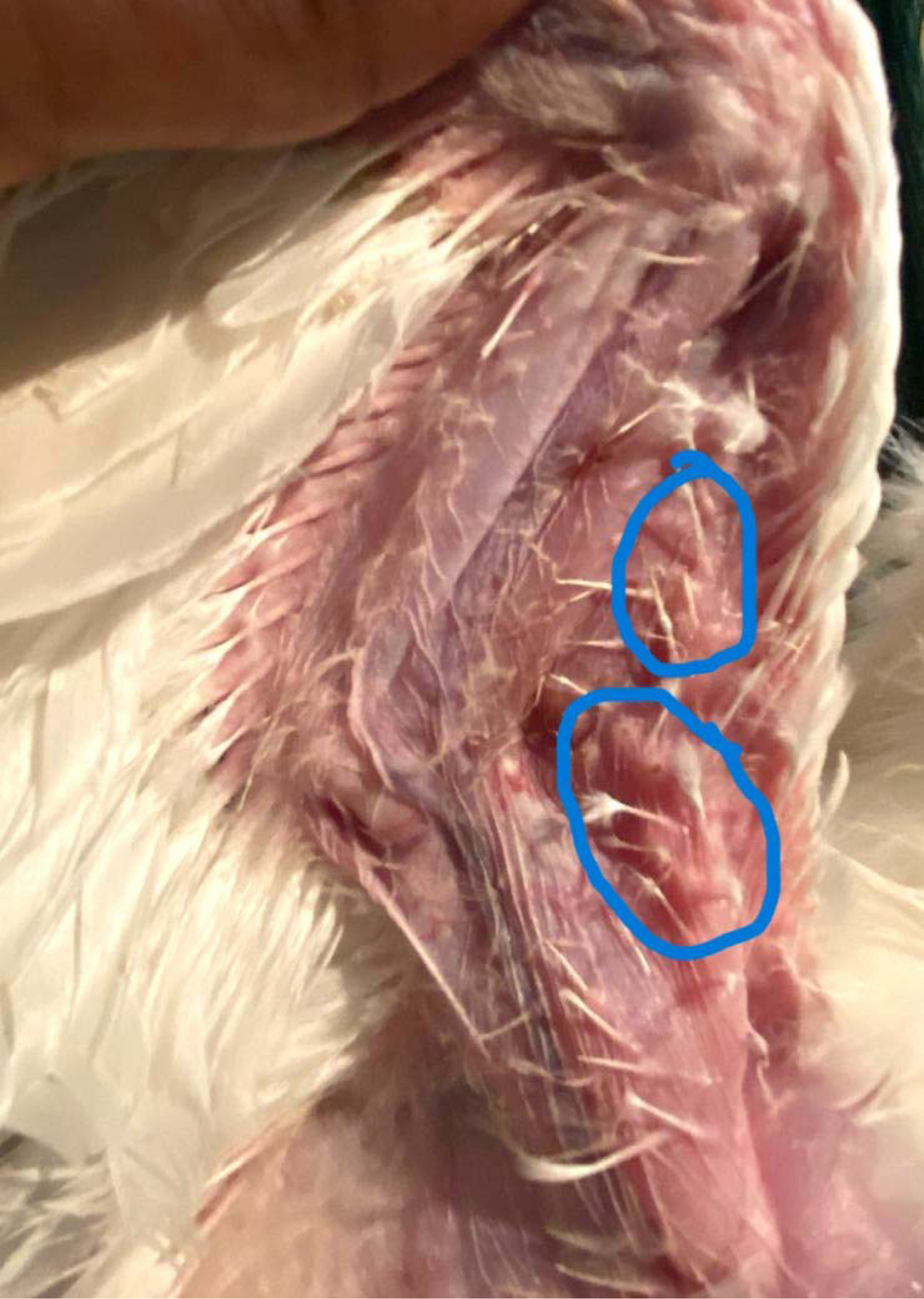
Vaccinated bird with CEF-adapted vaccine showing evidence of “takes” (Circled areas in blue).

## DISCUSSION

In this study, vaccine seed originally propagated on CAM were adapted to primary chicken embryo fibroblasts (CEF) cell culture. The adaptation of the vaccine virus to CEF cell culture in this study yielded positive results, producing CPE within 3-5 days, as reported in a similar study (Khalili *et al*., 2020). However, in this study, CPE was observed in some of the passages in less than 3 days (72 hours). The monolayer cultures in most of the passages were observed to have taken shorter time for attaining 80-90% CPE compared to the suspension cultures. This could be attributed to the advantage of the adsorption process in which the monolayer overlying medium is removed before the monolayer is inoculated with the suspension of viral organisms as opposed to the direct and immediate inoculation done during the suspension culture. The inoculation of the viral suspension onto the monolayer allows for wide surface contact between the virus and the cells. In addition, some studies have reported a delayed appearance of CPE with both FPV field isolate and vaccine strain. The CPE were only observed in some of the study at 3rd passage (Gilhare *et al*., 2015), while others at 8th passage (Khalili *et al*., 2020). However, in this study, CPE occurred at the first passage with a titer of 10^6.5^ TCID_50_/ml and 10^7.7^TCID_50_/ml for the BIOVAC and IZOVAC vaccine seeds respectively. This difference could be attributed to the nature and difference in the virus strain used in the cultures as it has been reported that not all strains can show CPE at the first passage (Khalili *et al*., 2020). In addition, a progressive increase in the titers with increased passages was observed which could be indicative of a successful adaptation of the virus to the CEF cell culture, as similar results were obtained in a study (Khalili *et al*., 2020).

The vaccination of the broiler chickens with various dilutions of CEF and CAM-based vaccines was also successful as “takes” were observed post-vaccination in all (100%) of the vaccinated birds, which is indicative of a successful vaccination. Similar results have been documented in several previous studies (Sarma and Sharma, 1988; Islam *et al*., 2008; Sarma *et al*., 2019). Generally, checking for post-vaccination “take” is considered one of the best methods for evaluating pox immunity in vaccinated chickens because properly administered efficacious vaccine is expected to show “takes” in 99-100% of the vaccinated chickens (Cookson, 1996; Barreda, 2016; Sarma *et al*., 2019). In addition, the over five weeks monitoring of the vaccinated birds, including groups that were vaccinated with ten times the standard field dose, showed no adverse reactions, death, or development of clinical signs of the disease as a result of the vaccine viruses. There was no reversion to virulence or any other safety issues in any of the vaccinated chickens vaccinated with different doses. These findings align with those of other reports or studies and agree with the recommendation of the OIE/WOAH (Shil *et al*., 2007; Sarma *et al*., 2019; Radwan and Mikhael, 2020; WOAH, 2023). Thus, the vaccines were found to be safe under experimental conditions as administering of the vaccine viruses at a much higher dose did not adversely affect the safety of the vaccines since all the vaccinated chickens remained healthy and were as active as the unvaccinated control birds.

Vaccination has remained the efficient means of control for the fowlpox virus in birds, and different vaccines are been used for this purpose, including vaccines developed from chicken embryo origin propagated on CAM (chorioallantoic membrane) and vaccines developed from cell culture origin adapted to avian origin cell culture. Certain studies have also been carried out to compare CAM-adapted and cell culture-adapted fowlpox vaccines, some of which have shown cell culture-adapted fowlpox vaccines to be better than the former (Baxi and Oberoi, 1999; Khalili *et al*., 2020). The finding of some studies also indicates that the cell culture-based vaccines performed better than the CAM derived vaccines. This was seen in the early observation of pox “takes” in vaccinated animals and the higher antibody titer as shown by ELISA results in birds vaccinated with cell culture-based vaccine as compared to the CAM-based vaccines. Plausible reason is the higher antigenic titer as a result of higher production rate and quality of CEF adapted viral vaccine compared to traditional CAM method of production.

To monitor the seroconversion of vaccinated birds with the different vaccines, ELISA and AGID was used to analyze the post-vaccination sera samples. Although, the ELISA result indicates seroconversion in the different groups, the baseline pre-vaccination sera samples indicated the presence of antibodies when analyzed with ELISA (giving OD values higher than the 0.216 critical cutoff) but were negative of antibodies when analyzed with AGID (indicated by the absence of precipitation lines in the test agar plates). However, the fact that the known negative controls were negative and the known positive controls were also positive validates the test. The discrepancy of the presence of antibodies in the baseline pre-vaccination sera samples could have been as a result of the presence of maternally-derived antibodies (MDA) in the chicks, even though the baseline pre-vaccination sera were sampled at 4 weeks of age, a period where MDA if present should have declined to negligible and non-protective levels (Islam *et al*., 2008). Generally, MDA is higher in day-old chicks and gradually declines below a positive level within 10-15 days of age (Natour *et al*., 1988; Akhter *et al*., 2008). This was however not the case in this study as the birds have high MDA at four weeks.

Amongst the various diagnostic techniques used for the detection of immune responses, AGID (Agar gel immunodiffusion) and ELISA (Enzyme-linked Immunosorbent Assay) are generally recommended for the determination of the immune status of individual animals or populations post-vaccination. However, despite the recommendations, these two diagnostic tests have also been reported to have their limitations (OIE Terrestrial Manual 2018). ELISA is considered to be a non-species-specific test (Buscaglia *et al*., 1985); however, some studies have shown ELISA to be highly sensitive but less specific when compared to AGID (Mockett *et al*., 1987; Buscaglia *et al*., 1985). In another study in which AGID was used to determine the seroprevalence of fowlpox antibodies in an area (Meseko *et al*., 2012), the resulting prevalence figures were found to be lower, and did not agree with a previous study which reported higher prevalence figures using ELISA technique (Ohore *et al*., 2007). This discrepancy associated with ELISA was attributed to the high sensitivity and low specificity of the test which makes it prone to false positive results (Buscaglia *et al*., 1985; Meseko *et al*., 2012). This discrepancy also occurred in this study with all the samples analyzed with ELISA showing the presence of antibodies, whereas the same samples run with AGID only showed 29% positivity to fowlpox antibodies. The baseline pre-vaccination sera samples that indicated the presence of fowlpox specific antibodies when analyzed with ELISA, were however negative or showed the absence of fowlpox specific antibodies when analyzed with AGID because though ELISA has higher sensitivity it can also sometime be less specific compared to AGID. Although the presence of MDA to fowlpox in the chicks has been ascribed to these findings, another plausible reason could have been a false positive result emanating from the non-specificity of the ELISA assay. Nevertheless, the AGID analysis showed progressive positivity after the first week post-vaccination, with a higher positivity observed in the fourth- and fifth-week post-vaccination.

There was also a higher positivity percentage with the group vaccinated with a standard dose of CEF-adapted fowlpox vaccine compared to the other vaccinated groups. The ELISA analysis also showed higher antibody titers (represented by higher optical density values) amongst the CAM-adapted and CEF-adapted vaccinated groups in the fourth- and fifth-week post-vaccination for the standard field and 10x doses which agrees with a finding which demonstrated an increasing titer of virus harvest with increasing passage for both CAM and CEF passages (Verma *et al*., 2015). This generally agrees with the AGID result obtained in this study, and also agrees with the report of other studies who also showed that fowlpox antibody levels significantly increase up to four weeks post-vaccination before it declines (Sarma and Sharma, 1988; Saini *et al*., 1990). In addition, chickens vaccinated with CEF adapted vaccine generally have higher OD values signifying the presence of higher antibody titer which aligns with the findings of some studies where birds vaccinated with the cell culture-adapted vaccine showed better protection in contrast to birds vaccinated with chicken embryo-adapted vaccines or where cell culture adapted vaccines were shown to be more effective (Siccardi, 1975; Baxi and Oberoi, 1999). In fact, cell culture-adapted (CEF) vaccine has been reported to have higher biological properties than a conventional vaccine prepared on the chorioallantoic membrane of embryonated eggs (El-Zein *et al*., 1974; Olfat *et al*., 2005), and are thus likely to provide more protection than the conventional CAM derived fowlpox vaccine, but both the CAM-adapted and CEF-adapted vaccines would elicit adequate immune responses (Baxi and Oberoi, 1999) as Fowlpox vaccine produced in embryonic chickens have the disadvantage of low level of infectious activity in chickens. In view of this, the search for fowl pox virus vaccine adapted to a cell culture with high infectious and immunogenic properties becomes critical (Yusifova, 2018; 2021).

Although most commercially available live Fowlpox vaccines are produced either by inoculation onto chorioallantoic membranes (CAM) of 9- to 12-day-old developing chicken embryos, the use of avian cell cultures such as chicken embryo fibroblasts (CEF), chicken embryo kidney cells, chicken embryo dermis cells, or the permanent quail cell line QT-35 are also some of the cell culture methods that have been used to propagate the fowlpox virus (Schnitzlein *et al*., 1988). In modern veterinary medicine, vaccines based on cell cultures have found widespread use (Yusifova, 2021; (WOAH, 2023). Fowl pox vaccine is also been manufactured using chicken embryo fibroblast (CEF) culture in many countries (Khalili *et al*., 2020), and cultivation of fowl pox virus in chicken embryo cell cultures is common in preparing biomass for vaccine production (Yusifova, 2018). Thus, the use of cell culture-adapted vaccines for fowlpox has been reported to be more economical and productive compared to CAM-adapted vaccines whose production is more time-consuming, thus many manufacturers began to produce vaccines based on cell systems (Yusifova, 2021). Thus, the use of different cell culture and different strain of viruses by researchers to improve and increase vaccine effectiveness has been ongoing (Yusifova, 2017; 2021). The need therefore to develop more potent vaccines in enough quantity to meet the demand of the Nigerian poultry is very important.

## CONCLUSION

In conclusion, this research successfully demonstrated the development and adaptation of Fowlpox virus to a primary cell line of CEF for the propagation and titration of a vaccine virus. The study involved multiple passages in both suspension and monolayer cultures, resulting in effective virus adaptation to CEF for vaccine production, with notable cytopathic effects observed. The vaccine virus, propagated in both CAM and CEF cultures, exhibited high titers, indicating successful production. Immunogenicity and safety assessments of CEF-adapted and chorioallantoic membrane (CAM) vaccines in broiler chicks revealed the induction of “takes” without adverse effects, confirming the vaccines’ safety. Vaccines produced in CEF were immunogenic, safe, and produced no untoward clinical signs or adverse effects in vaccinated chickens. Overall, these findings support the safety, and immunogenic potential of the developed CEF-adapted vaccine, laying a solid foundation for its potential application in controlling fowl pox virus infections in poultry. The application and production of fowlpox vaccine by cell culture can be a small step towards the self-sufficiency of this type of vaccine in Nigeria. Vaccination remains the most effective means of controlling and preventing fowlpox disease in poultry. From the results obtained in this study, it becomes imperative that further studies be carried out on a larger scale to establish the efficacy, safety, and potency of CEF-adapted fowlpox vaccines.

## Acknowledgements

The authors acknowledge the African Union Commission for awarding the grant for this research through the Pan African University Life and Earth Sciences Institute (including Health and Agriculture), Ibadan, Nigeria. We also acknowledge the Viral Vaccine Production Unit of the National Veterinary Research Institute (NVRI), Nigeria for the laboratory support and the Regional Laboratory for Animal Influenza and Transboundary Animal Diseases NVRI for the serology testing.

